# Improving genome-scale metabolic models of incomplete genomes with deep learning

**DOI:** 10.1101/2023.07.10.548314

**Authors:** Meine D. Boer, Chrats Melkonian, Haris Zafeiropoulos, Andreas F. Haas, Daniel Garza, Bas E. Dutilh

## Abstract

Deciphering the metabolism of microbial species is crucial for understanding their function within complex ecosystems. Genome-scale metabolic models (GSMMs), which predict metabolic traits based on the enzymes encoded in a genome, are promising tools to study microbial ecosystems when genome sequences can be obtained. However, constructing GSMMs for uncultured bacteria is challenging, as metagenome-assembled genomes are typically incomplete, leading to metabolic reconstructions with numerous gaps. Existing methodologies often fill these gaps with the minimum set of reactions necessary to simulate an objective function such as growth. Here we introduce an artificial intelligence-based alternative: the Deep Neural Network Guided Imputation Of Reactomes (DNNGIOR). The DNNGIOR neural network learns weights for missing reactions in incomplete GSMMs from patterns in the presence and absence of metabolic reactions in genomes spanning the bacterial domain. We identified two important factors contributing to prediction accuracy: (1) the frequency of reaction across all bacteria, and (2) the phylogenetic distance between the query and the genomes in the training dataset. Reactions that occur in > 30% of the training genomes can be accurately predicted (Mean F1 score = 0.85). The weights generated by the DNNGIOR network improved the gap-filling of incomplete GSMMs, when assessed on a large and phylogenetically diverse testing dataset and a small set of high-quality manually curated models. The accuracy of DNNGIOR was on average 14 times greater than the standard unweighted gap-filling for draft reconstructions, and 2-9 times greater for manually curated models. DNNGIOR models could also simulate experimentally measured carbon usage profiles with similar accuracy as CarveMe. DNNGIOR is available at https://github.com/MGXlab/DNNGIOR or as a pip package (https://pypi.org/project/dnngior/).

## Introduction

Simulating microbial metabolism is an effective method to understand bacterial physiology and interactions within their communities(1–3). The functions and interactions of bacteria can be inferred from their genome sequences using genome-scale metabolic models (GSMMs)(3–6). GSMMs can be constructed either manually or automatically with tools such as RAVEN (7), ModelSEED (8), KBase (9), and CarveMe (10), which identify metabolic reactions encoded on the genome and build a metabolic network. However, if the original genome sequence is incomplete, a common occurrence with meta-genome-assembled genomes (MAGs), the inferred GSMM will also be incomplete (11). Consequently, gaps in GSMMs emerge due to missing knowledge and errors introduced during sequencing (12), binning (13,14), and annotation (15). In the past, gap-filling was primarily executed through manual curation (16–18), but this method is time consuming and does not scale well for studies that include a large number of GSMMs (19–21).

Several algorithms have been developed to automate gap-filling, such as FastGapfilling (22), GlobalFit (23), CHESHIRE (24) and OptFill (25) that add reactions that allow a GSMM to simulate growth or match phenotypic profiles. The reaction sets that can gap-fill a model are not unique (26) and the organism’s actual metabolism may not always align with the minimal set of reactions satisfying a user-defined objective (27). This indicates room for refining gap-filling algorithms to yield more realistic solutions. As most gap-filling algorithms allow us to weigh reactions individually according to their likelihood of being in the model (10,22,26), several attempts have been made to find weights based on genomics (28), proteomics (29), topology (26,30) or reaction type (8). Nevertheless, despite these advances, determining the optimal weights for any reaction and any model still remains challenging (4).

In this article, we introduce DNNGIOR: a python package that uses a neural network to assign weights to metabolic reactions to complete GSMMs that are built from incomplete genomes. This neural network is trained to discern patterns in the cooccurrences of reactions across the bacterial domain and to predict reactions based on incomplete reaction sets, with the goal of assessing which reactions may be missing from an incomplete network. This information will be useful for automated and manual GSMM reconstruction. When we used the predictions of the neural network to weight reactions in incomplete GSMMs, we found that the accuracy of the neural network depends on the frequency of reactions in the training data and the phylogenetic relatedness of genomes used to generate this data. We benchmarked the predictions using both automated and manually curated models, including data from a recent study on carbon usage profiles by plant-associated bacteria (31).

## Methods

### Collection and processing of the training and testing data

We constructed the training and testing datasets using genomes collected from the BV-BRC database ((32), formerly PATRIC, accessed 26th April 2022). For training, one genome per species was selected based on sequencing quality scores using the formula: completeness - (5 * contamination). Ties for this score were resolved by selecting the genome with the highest coarse consistency. This selection resulted in a dataset of 13,359 genomes (Supplementary Table 1) that comprehensively represents the bacterial domain while reducing the risk of overfitting on well-studied species. From this dataset we set aside one best genome from each of the 1,659 genera based on the same score, resulting in a training dataset of 11,700 genomes and a testing dataset of 1,659 genomes (Supplementary Table 2). This ensured that the testing dataset contained diverse bacterial genomes, not biassed towards genera with more species that were different from those in the training dataset.

For the 1,659 genomes in the testing dataset, we created a phylogenetic tree using concatenated alignment of hits to HMM profiles of 71 single-copy marker genes that was used for visualising and further investigating the performance of our approach. Phylogenetic distances in the tree represent the number of amino acid substitutions per site.

From all genomes, metabolic models were constructed using either ModelSEED (8) or CarveMe (10) and the set of gene-associated reactions was determined. From these models we determined the total set of reactions that were annotated in the 13,359 genomes, resulting in a “pan-reactome” of the bacterial domain within the ModelSEED and BiGG databases (n = 2543 and 4240 reactions, respectively). These pan-reactomes contain all reactions for which predictions can be made. Most of the other reactions in the ModelSEED and BiGG databases either originate from non-bacterial organisms (e.g. plants, fungi, animals) or are artificial reactions. Both these categories of reactions are not associated with bacterial genes, and therefore do not appear in the draft models or associated reactome. We decided against including non-gene-associated reactions in the training data to avoid learning the biases that automated tools introduce when including non-gene associated reactions. For every model in the training and testing datasets, we constructed a binary array describing which reactions were present, resulting in an incidence matrix of all reactions in all genomes.

During training we repeatedly deleted 30% (*<u>n</u>* ≈ 300) of the reactions in each genome, this was done 30 times for each of the 10,700 genomes resulting in a training dataset of 351,000 incomplete reaction sets. Reactions were randomly deleted either according to a uniform probability (Figure 1a) or with a bias towards lower frequency reactions (Figure 1b).

**Figure 1:**
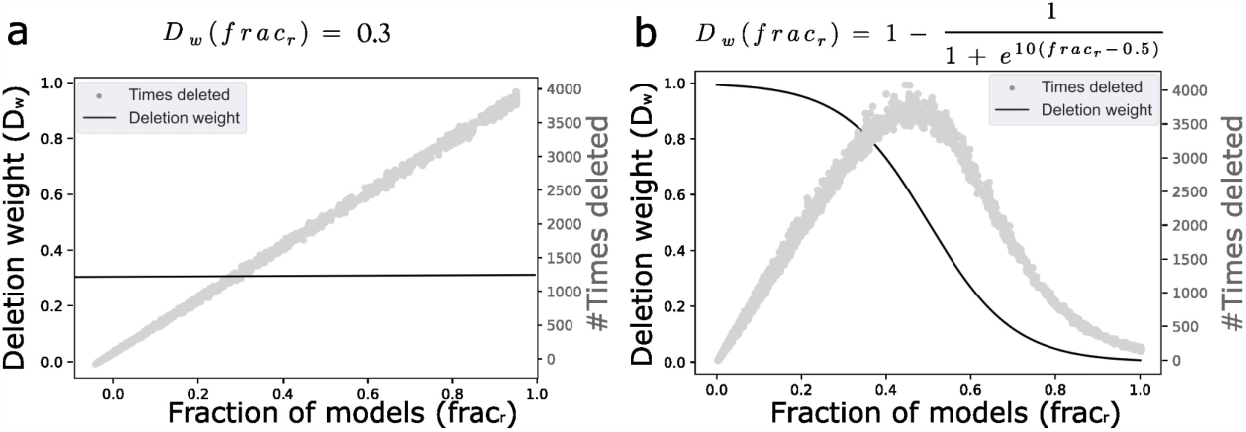
Probability function for uniform (a) or weighted (b) deletion of reactions in the training and testing datasets. Based on these weights, the scatter plots show how often reactions were actually deleted on average per replicate. ***Frac***_***r***_ represents the fraction of genomes in the training dataset that include a reaction.

We also generated several additional training datasets with certain phyla purposefully excluded to explore the importance of a full representation of the bacterial domain. For each genome, the original reaction sets as predicted by the genome-based draft reconstructions were used as truth. Testing datasets were created in a similar manner.

### Hyper-parameterization and loss function of the neural network

Two neural networks were built, one for predicting ModelSEED reactions and one for predicting CarveMe or BiGG reactions. Both neural networks were built using Tensorflow(33) v2.0.0. Their topology consists of an input and output layer of 2,453 or 4,240 nodes (one for each reaction in the ModelSEED or CarveMe pan-reactomes respectively) and three hidden layers. All layers were fully connected resulting in a network with 1,260,697 or 2,306,960 parameters respectively. The optimizer used for training both networks was the Adam optimizer (34) with the following parameters: learning rate = 0.005, beta1 = 0.9, beta2 = 0.999, epsilon = 1.0e-8, decay = 0.01. Hyper-parameterisation was performed in 100 trials using the Optuna package (35) v2.0, resulting in the following hyper-parameters: number of nodes per hidden layer = 256, batch size = 50, number of hidden layers = 3, dropout = 0.1, number of epochs = 10.

For the loss function, we used a customised version of the binary cross-entropy function (CE, Equation 1).

**Equation 1:** Loss function based on binary cross-entropy (34) to calculate the difference between the neural network output 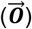 and the correct reactions 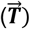. 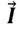 is the input vector of the NN, b0 is the absent class scaling factor.

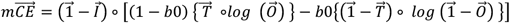

CE is calculated as the log-loss of the difference between what the network predicts 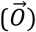 and the truth 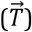. We introduced two adaptations to increase the performance based on our training data. First, as for a given reaction set only ∼960 of the possible 2,457 reactions are present, we introduced a scaling factor (*b*_0_ = 0.3) that allowed us to scale the loss of the two classes (absent and present). We multiplied *b*_0_ by the loss for the absent class and 1 – *b*_0_ by the loss for the present class. Second, we added a masking vector (1-T) that allowed us to exclude the loss associated with predictions for reactions that were already known to be part of the genome, i.e. those given as input to the neural network. We multiplied the loss of both classes element-wise 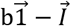 where 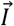 is the vector of reactions given as input. This adaptation ensures that the neural network learns to complete the reaction set and does not simply repeat the input.

### Gap-filling algorithm and database

After predicting weights for all reactions, we used those weights to guide a half-interval search for the minimal set of reactions that simultaneously has a high probability and generates biomass flux that is greater than zero. The half-interval gap-filling algorithm was adapted from (22). Briefly, this algorithm iteratively minimises the following objective function with linear programming conditional on flux through the biomass reaction (f_b_):

**Equation 2:** Objective function of the half-interval gap-filling algorithm that optimises gap-filling of an incomplete metabolic network based on weights and flux of reactions in a network.

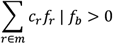

This sums over all reactions (r) in the candidate set (M), with f_r_ the flux through reaction r and c_r_ a user-defined cost used to implement the different weighting schemes during gap-filling. By using linear programming, the runtime is reduced by up to two orders of magnitude compared to mixed integer linear programming (22).

The reaction database from where reactions were selected was downloaded from the BiGG website (http://BiGG.ucsd.edu, Supplementary Table 3) for the BiGG and CarveMe models and from the ModelSEED website (https://modelseed.org, Supplementary Table 4) for the ModelSEED models. From these databases, biomass reactions were removed. Reversible reactions were split into two reactions, one for each direction. The algorithm can also take into account different media compositions, as these may affect the solution.

### Weighting schemes for guiding the gap-filling algorithm

Four weighting schemes were used to guide the half-interval algorithm, namely: W1. No weights, W2. Naive binary weights, W3. Frequency weights and W4. NN weights (see Equation 3). For W1, all reactions in the database (n = 43,775) are given the default cost (c_r_ = 50). For W2, a low fixed cost (c_r_ = 1) is given to reactions that are present at least once in the training dataset (*R*_*train*_, n = 2457), and the default cost to all other reactions in the database. For W3 a lower cost is given to a reaction if it is present in a higher fraction of genomes in the training dataset (Equation 3). For W4, a lower cost is given to reactions that have a higher prediction score. Because the half-interval gap-filling algorithm tries to minimise the product of a reaction’s cost and flux, reactions that are given lower costs are given more weight in the final solution.

**Equation 3:** Four weighting schemes to guide the half-interval gap-filling algorithm, see text for details.

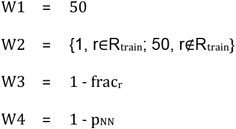

We compared the four different weighting schemes by gap-filling the models that were constructed based on the 1,659 genomes from the testing dataset from which 30% of reactions (*<u>n</u>* ≈ 300) was deleted at random in triplicate. After gap-filling, we counted which removed reactions were re-added correctly (TPs) or not (FNs), and which reactions were falsely added (FPs). These were used to calculate a F1-score for the different weighting schemes.

### Curated genome-scale metabolic models

We selected six high-quality manually curated models based on literature (iML1515 (16), iJB785 (36), iYL1228 (37), iND750 (38), iYO844 (39), and iYS854 (40) from the BiGG database (41). We deleted in 10x replicate 30% of reactions from these models and gap-filled them with the same four weighting schemes as for gap-filling the draft-models (see section above). For these models the neural network trained on CarveMe models was used as the reaction identifiers matched those from the curated models. To illustrate which reactions are most likely to be found with the different weighting schemes we also deleted in 500x replicate 30% of reactions from the *Escherichia coli* model (iML1515) and constructed an Escher map (42) of the central metabolism.

### Validation of gap-filled models based on experimental data

We obtained the genome sequences of 224 bacterial isolates from Arabidopsis thaliana leaves (31). Draft models were constructed using ModelSEEDpy (43) v0.3.0 and gap-filled in a minimal medium (Supplementary table 5) with and without DNNGIOR neural network weights. CarveMe models were built using version v1.6.0 with default parameters and the M9 minimal medium provided by the package. Simulated carbon utilisation profiles were established by measuring flux through the biomass function (‘bio1’ for DNNGIOR models and ‘Growth’ for CarveMe models) on 45 different carbon sources using cobrapy v0.28.0. Balanced accuracy scores were calculated by comparing the simulated carbon utilisation profiles to those that were measured in vitro (31).

**Equation 4: Balanced accuracy score**.

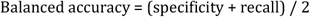

## Results and Discussion

We set out to build a tool that would make use of the co-occurrence patterns of reactions found in a broad range of bacterial genomes to improve the reconstruction of metabolic models from incomplete genomes and MAGs. Therefore, we trained a neural network on the occurrence of 2,457 or 4,240 metabolic reactions for ModelSEED and CarveMe, respectively, in over 13 thousand species (training and testing). This network predicts missing reactions within incomplete reaction sets. A schematic overview of our approach is depicted in Figure 2.

**Figure 2:**
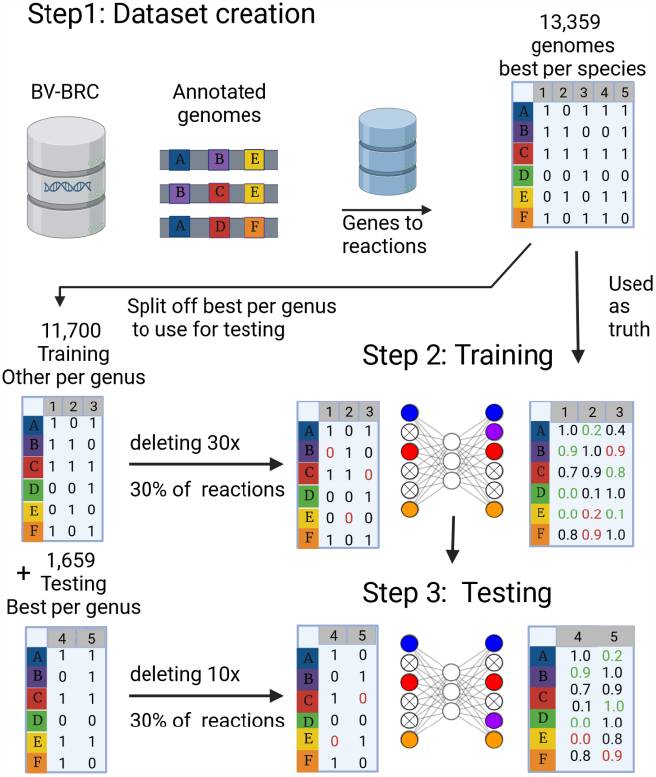
Schematic overview of training and testing the DNNGIOR neural network. Step 1: Constructing the dataset. Genomes were collected from the BV-BRC (32) database selecting one genome per species (13,359 genomes), genomes were annotated and metabolic networks constructed as outlined in Methods section. The resulting incidence matrix of reactions in different genomes was split into subsets for testing (best per genus, 1,659 genomes) and training (remaining 11,700 genomes). Step 2: Training the neural network. We randomly deleted 30% of reactions 30x to simulate incomplete genomes. The network was trained to predict which reactions were removed, while not predicting the reactions that were not part of the original draft model. All reactions that were given as input to the network are ignored when calculating the loss from the predictions. Step 3: Testing the neural network. To estimate the prediction accuracy for different reactions in a diverse set of organisms, the network was tested on the 1,659 genomes in the testing set. Incomplete genomes were simulated as explained above, 10x per genome. This figure was created using BioRender (44).

### Reaction frequency is an important factor for prediction accuracy

Before testing the predictions to guide gap-filling we first wanted to understand the factors underlying accurate predictions by the DNNGIOR neural network. Understanding these factors can show the strengths of the network, show possible areas of improvement, and provide new insights into the gap-filling problem. We identified two important factors that affect the accuracy of predictions by the neural network: (1) the frequency of reaction across all bacteria and (2) the phylogenetic distance between the organisms in the testing and training dataset.

Frequent reactions have higher recall and precision than rare reactions, with core reactions that are present in > 90% of all bacteria having a recall of 0.96 (sd = 0.007) and precision of 0.86 (sd = 0.039) (Figure 3a). The neural network has more opportunities to learn reactions and their associated patterns than when a reaction is only present in a few genomes.

**Figure 3:**
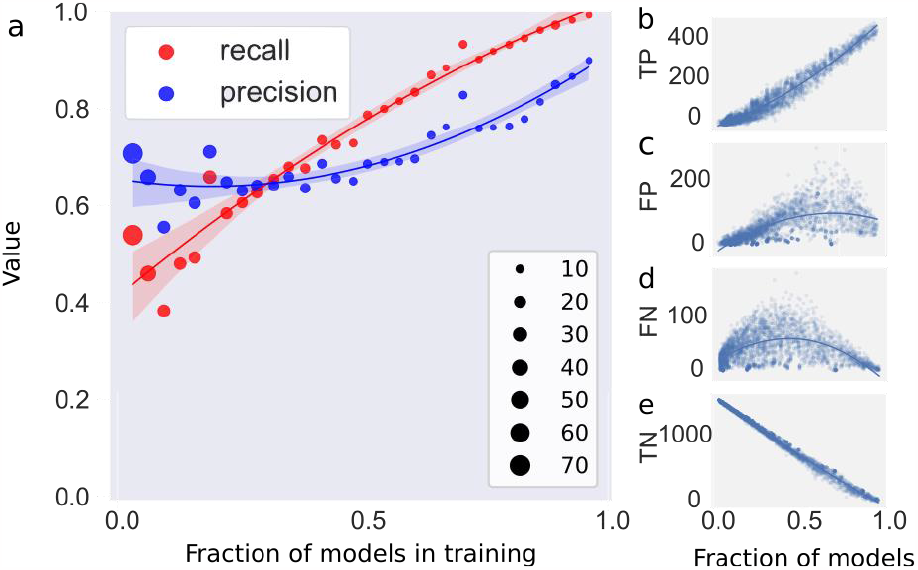
Prediction accuracy increases with reaction frequency. a) Relationship between the recall (red) and precision (blue) of predictions as a function of the reaction frequency. Reactions were binned in ranges of 50 reactions, dot size corresponds to bin size, on the y axis precision and recall as a fraction. The shaded region shows the 95% confidence interval of the regression. Recall = TP/(TP+FN), Precision = TP/(TP+FP) b-e) Regression plots of True Positives (TP, b), False Positives (FP, c), False Negatives (FN, d) and True Negatives (TN, e) as function of the fraction of genomes a reaction is present in. Trend lines were estimated using a Polynomial Regression Model.

Specifically, true positives (TPs) increase linearly with frequency (Figure 3b) while true negatives (TN) decrease linearly (Figure 3e). Since the total number of times a reaction is deleted also increases with frequency, there are more opportunities to correctly predict that a reaction is present or absent. Moreover, the false positives (FPs) and false negatives (FNs) decrease for more frequent reactions (Figure 3c and d). This indicates that the neural network can accurately predict whether a reaction should be present or absent from the model if they are sufficiently represented in the training set.

### Training data can be adapted to reflect the frequency of missing reactions in real data

As we have seen above, the frequently occurring reactions can be more accurately predicted. However, the most frequent reactions might not be the ones that are most likely to be missing from incomplete models in practice. Several biases in binning and annotation affect the distribution of missing reactions in real data. For instance, accessory genes are more likely to be missing from MAGs because the binning of mobile genetic elements is less effective than the rest of the genome (14). Additionally, large genomes with many accessory genes are more difficult to annotate than smaller genomes with mostly core genes (15). As accessory genes are by definition rarer than core genes (45), this leads to a bias towards rare genes being missing from MAGs. Another important bias is that taxa with less-researched members are more difficult to annotate accurately using homology-based tools (15,46). Finally, reactions can be perceived to be rarer because they are more often missing from the data.

These biases could be a concern if we want to use DNNGIOR to gap-fill models based on larger genomes, MAGs, or from the less researched parts of the bacterial domain as reactions missing from those models might likely be rare. Therefore, we tested the DNNGIOR neural network on data that contained a deliberate deletion bias, where rarer reactions were deleted more often (see Figure 1 in the Methods). In this case, the F1-score decreased 36% as the network overestimated common reactions and underestimated rare reactions (Figure S1). In contrast, if the network was trained on data with the same bias towards deleting rarer reactions, this effect was reduced (from 36 to 18%), and the network became better at predicting rare reactions (Figure S1). This means that it is possible to train the network to account for biases that could potentially be present in the data. However, this and the bias is difficult to quantify, the DNNGIOR package uses a neural network trained on random deletion by default.

### Short phylogenetic distances and complete representation improve prediction

We wanted to determine next which genomes are predicted better. We expected that predictions would be better for genomes from well-sampled taxa than from taxa with only few sequenced relatives. To test this, we plotted the F1-score for all genomes in the testing dataset on a phylogenetic tree (Figure 4a). While the DNNGIOR neural network scored well on most models (mean F1-score = 0.84, sd =0.054, Figure 4b), predictions were more accurate for species that had close relatives in the training data than for species that were more distantly related. When we made predictions for every species in the testing dataset, we found that the F1-score of the predictions correlates with the distance to the closest neighbour (Pearson r^2^ = 0.261, p = 1.53*10^-40^). Indeed, species that have close relatives in the training data are easier to predict than species that are phylogenetically unique (Figure 4c).

**Figure 4:**
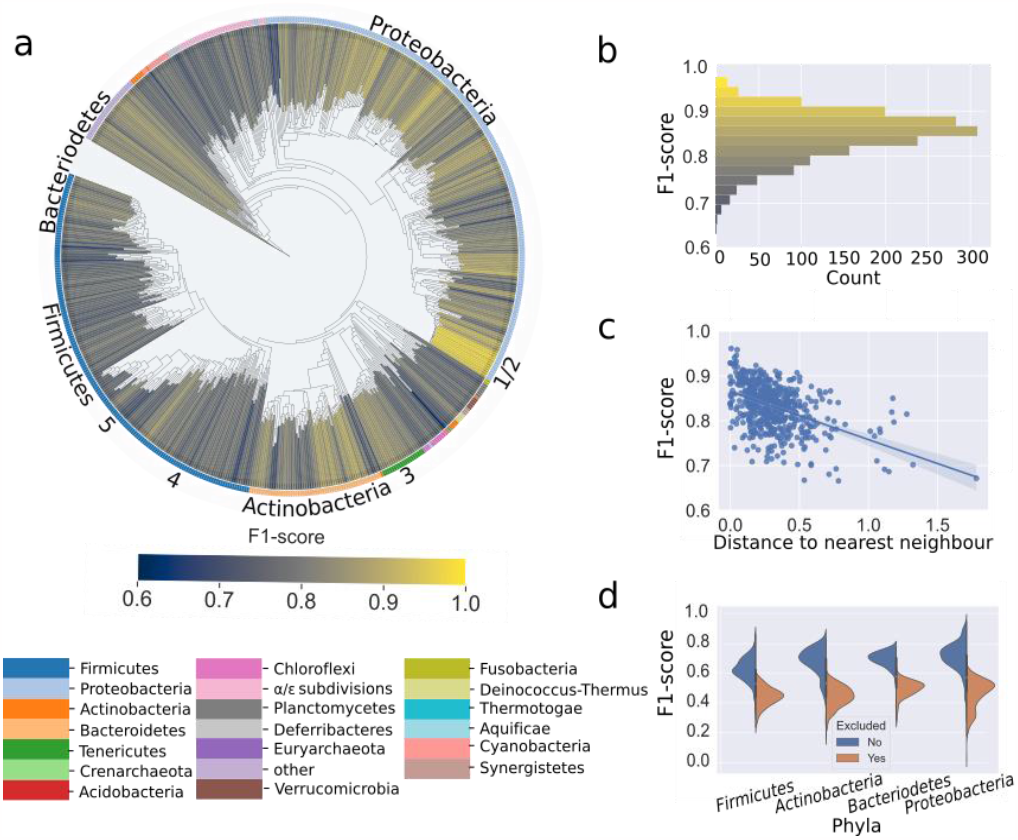
Phylogenetic distance influences the accuracy of DNNGIOR neural network predictions. a) Phylogenetic tree based on a concatenated multiple sequence alignment of 71 single-copy marker genes of all the 1,659 genomes in the testing dataset. Branch colour represents the F1-score, the colour of the outer ring corresponds to the phyla ordered by size, all phyla with less than five species are combined in other, the four largest phyla are annotated. The five species with high-quality curated metabolic models used during the gap-fill analysis are also marked: 1. Escherichia coli, 2. Klebsilla pneumoniae, 3. Synechococcus elongatus, 4. Bacillus subtilis, and 5. Streptococcus aureus. The tree is based on a concatenated alignment of hits to HMM profiles of 71 single-copy marker genes. Phylogenetic distances in the tree represent the number of amino acid substitutions per site. b) Histogram of the F1-scores coloured with the same colour map as the Tree. c) Scatterplot of the F1-score versus the distance to the nearest neighbour expressed in the Jaccard distance, trendline (Pearson r2 = -0.51, p = 1.53*10-40). d) Split violin plots of F1-scores of the predictions for neural networks when different phyla are included (left) or excluded (right) from the training data.

The correlation between F1-score and interspecies distance illustrates the importance of a good and complete representation of the bacterial domain as this would reduce the average distance between query species and those in the training data. We confirmed this hypothesis by excluding some phyla from the training data, which resulted in lower F1-scores compared to when all phyla were included (Figure 4d). We also confirmed that the reverse is true, i.e. training on only reaction sets from one phylum increases the performance of that phylum (Figure S2). Using DNNGIOR specialised neural networks can be trained exclusively on genomes from a certain phylum or biome. This sacrifices general applicability, but it may be expected to yield a (moderate) increase in performance. However, as biome annotations are often subjective and may be either redundant or ambiguous (47), we opted to train a single model and present DNNGIOR as a universal tool that can be directly applied to genomes and MAGs from all phyla and derived from any biome.

### Neural network weights improve gap-filling of draft models

Next, we determined the effectiveness of using NN-predicted weights to guide gap-filling, compared to other alternatives. For this, we gap-filled the models from the test dataset using the half-interval gap-filling algorithm (see Methods), where weights may be assigned to individual reactions and the algorithm finds a metabolic network that is capable of generating a biomass flux, while minimising the weights that are added overall. We assessed whether the artificially removed reactions could be recovered when reactions were weighted based on four different weighting schemes: W1. No weights, W2. Naive binary weights, W3. Frequency-based weights, and W4. NN weights (see Methods). To be clear, the process of gap-filling results in a functional model that can produce biomass, while the training set consists of draft models that generally cannot produce biomass and contain exclusively reactions derived from genome annotations, from which a fraction was deleted (see Figure 2). We found that NN weights (W4) significantly improved the accuracy of automated gap-filling compared to other weighting schemes (Supplementary Table 9), i.e. they allowed a larger fraction of the deleted reactions to be recovered in the model. By using scheme W4, the F1-score increased 13.98 times compared to W1 (p = 4*10^-18^), 1.92 times compared to W2 (p = 4*10^-18^) and 1.09 times compared to W3 (p = 3*10^-18^, Figure 5).

**Figure 5:**
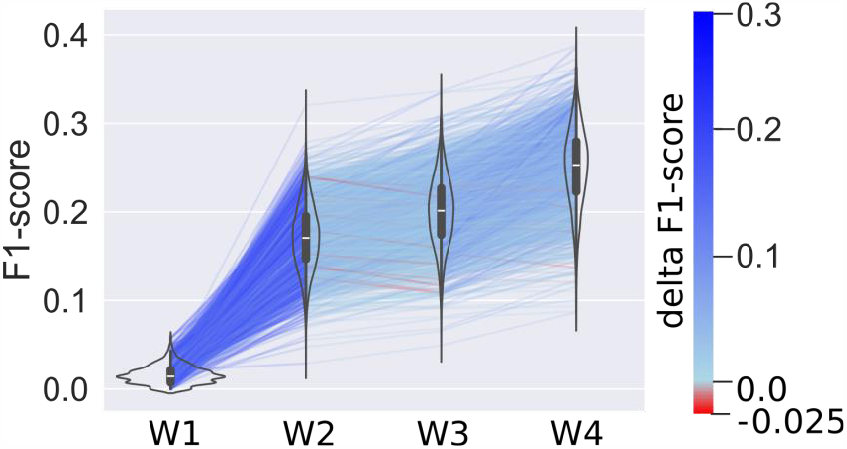
Weighted gap-filling of draft models. Violin plots of F1-scores of the gap-filling of 1,659 models in the testing set, from which we randomly deleted 30% of reactions in triplicate. These reduced models were gap-filled using four different weighting schemes (equation 3). For W1 (“No weights”) all reactions in the database are weighted equally. For W2 (“Naive binary weights”) all reactions that are present in the training data were given the same weights. For W3 (“Frequency weights”) the frequency of the reaction was used to weigh reactions (see Equation 3). For W4 (“NN weights”) the prediction scores generated by the DNNGIOR neural network were used. Lines connect the same models to show trends and are coloured based on the difference in F1-score for the model between weighting schemes. All groups were significantly different (Supplementary Table 8).

When we try to explain the improvement observed, a major part is already visible with W2 which shows that the reactions from the pan-reactome are indeed the most important ones. The improvement with W3 is in line with the observations from assessing the accuracy of the predictions of the DNNGIOR neural network directly, namely that frequency of a reaction is also important for gap-filling. Although this may be expected, the reaction frequency has often been neglected when gap-filling strategies are developed. These strategies often focus on flux or network topology, giving the same cost to all reactions. This leads to addition of a minimum number of reactions that are necessary for growth, agnostic to all external information. Here, we found that simply weighing reactions by their frequencies in the bacterial domain already significantly improves the accuracy of gap-filling (Figure 5). The DNNGIOR neural network scores (W4) further improve gap-filling, suggesting that additional information has been learned from the co-occurrence patterns.

When observing these results, we note that the F1-scores were lower than might be expected based on the prediction accuracy (Figure 3b). This consistent trend can be attributed to the fact that the draft models were already incomplete before the additional reactions were removed, because they were based on genome annotations alone. Thus, some reactions were necessarily added by the half-interval gap-filling algorithm to enable biomass production that were counted as false positives here since they were not present in the genome annotation. Furthermore, the objective of the half-interval gap-filling algorithm was not to find back reactions but rather to find a set of reactions that allows biomass production, while minimising their overall weights. As many of the annotated reactions that were removed were not strictly required for biomass production they were not added back, leading to false negatives. Although the F1-scores were thus systematically reduced, we can still interpret the trends in performance of the four different weighting schemes in Figure 5.

Overall, reactions that were assigned a high probability by the DNNGIOR neural network are likely to be present in the metabolic network, whereas low probability reactions are likely to be absent. To further incentivise the inclusion of high probability reactions while still satisfying the biomass production objective, we provided these reactions with negative, rather than zero costs (see Equation 2). These reactions were thus even more stimulated to be included, and indeed this nearly doubled (x1.95) the F1-score compared to using only positive weights (Figure S3a). However, this approach also led to an increase in false positives for reactions that were absent but were still assigned high probabilities by DNNGIOR (Figure S3b) (46).

### Neural network-based weights improve gap-filling of curated models

Next, we assessed the performance of the four weighting schemes using six high-quality manually curated models. As above, we artificially removed reactions from these models and tested how well these reactions were reintroduced when weighted by the DNNGIOR neural network (see Methods). As shown in Figure 6, the NN weights (W4) outperformed the other weights for all six models. Notably, *Saccharomyces cerivisiae* (iND750) performed the worst of the six tested models (mean = 0.11, sd = 0.05). This may be expected since *S. cerevisiae* is a eukaryote and only distantly related to the prokaryotic reference genomes that comprised the DNNGIOR training dataset. The best performance (mean = 0.22, sd = 0.08) was found for *Escherichia coli* (iML1515), a species from a well-studied family with many reference genomes. The good performance of the E. coli model, but also of the other models might in part be explained by the fact that they are derived from relatively well-studied taxa with extensive annotation. Thus, these models perform better than most draft models in our testing dataset above.

**Figure 6:**
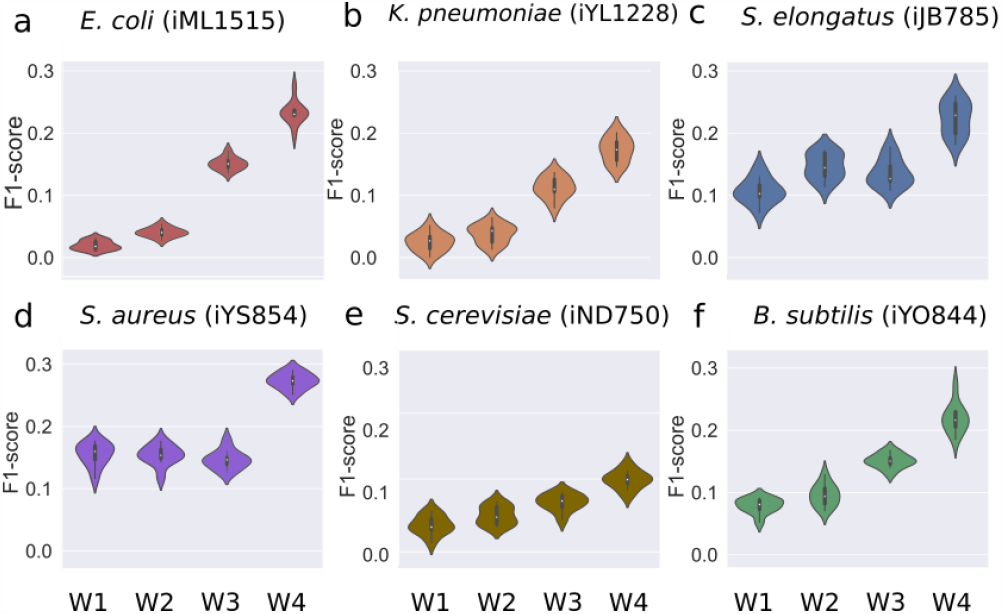
Weighted gap-filling of six curated metabolic models. Violin plots of F1-scores of the gap-filling of curated models, from which we randomly deleted 30% of reactions 10 times. These reduced models were gap-filled using four different weighting schemes (Equation 3). For W1 (“No weights”) all reactions in the database are weighted equally. For W2 (“Naive binary weights”) all reactions that are present in the training data were given the same weights. For W3 (“Frequency weights”) the frequency of the reaction was used to weigh reactions. For W4 (“NN weights”) the prediction scores generated by the DNNGIOR neural network were used. a) E. coli iML1515, b) K. pneumoniae iYL1228, c) S. elongatus iJB785, d) S. aureus iYS854, e) S. cerevisiae iND750, and f) B. subtilis iYO844.

The fact that the NN weights (W4) performed better than the Frequency weights (W3) indicates that the neural network learned more than the reaction frequency alone. To gain an intuition for the kind of additional information the network could have learned, we built Escher maps of the citric acid cycle of *E. coli* and coloured reactions by their mean recall over 500 iterations where we randomly deleted 30% of the reactions each time (Figure 7). Scheme W4 produced the highest recall (Figure 7d), followed by W3(Figure 7c). We found that recall partially correlated with the reaction frequency in the training data for both W3 (Pearson r^2^ = 0.5, p = 1.06*10^-116^) and W4(Pearson r^2^ = 0.25, p = 1.22*10^-26^, Supplementary Figure S4b) but that W4also found some rare reactions that were more specific to *E. coli*, indicating that the network also learned specific co-occurrence patterns. The rest of the central metabolism of *E. coli* showed a similar pattern as the citric acid cycle (Supplementary Figures S5 and S6).

**Figure 7:**
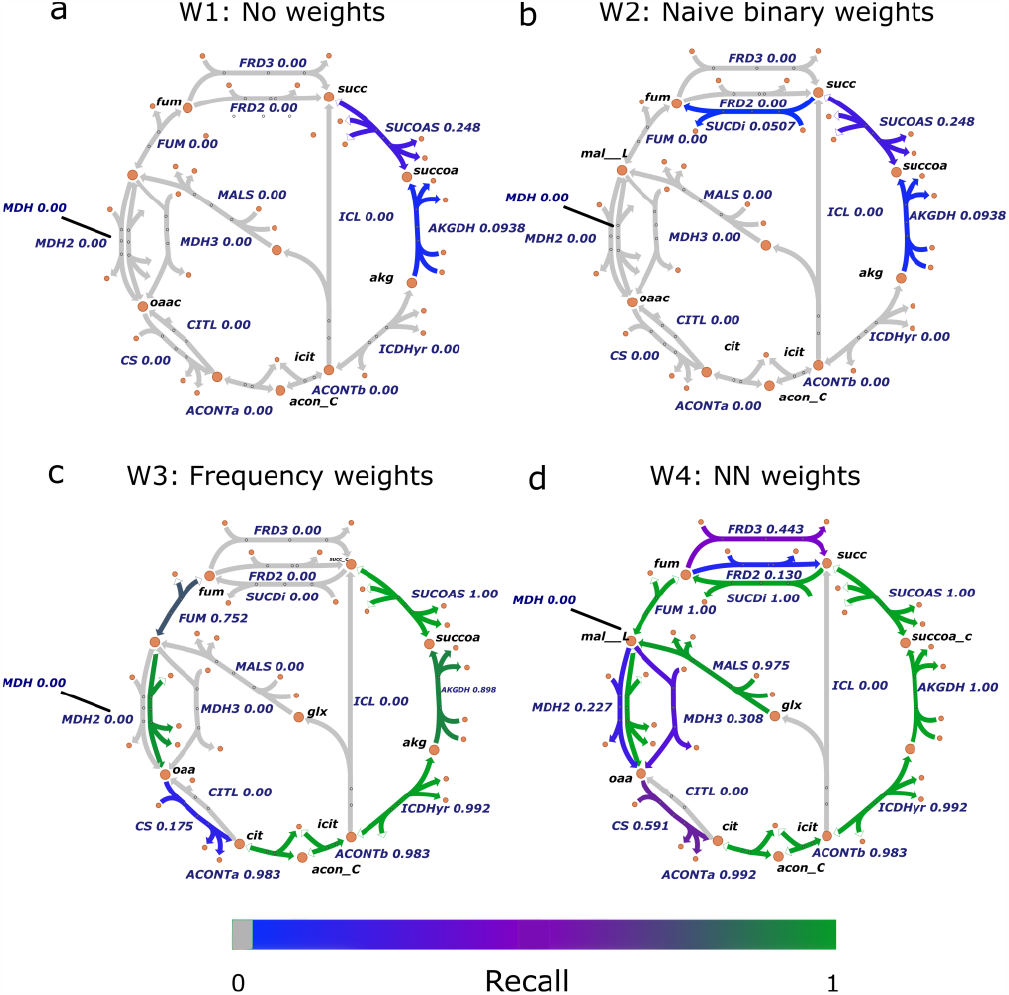
Recall of the reactions in the citric acid cycle of *E. coli*. Escher maps of the citric acid cycle coloured by the recall after gap-filling models, from which we randomly deleted 30% of the reactions 500 times. Reduced models were gap-filled using four different weighting schemes: a) W1: No weights b) W2: Naive binary weights c) W3: Frequency weights and d) W4: NN weights (see Methods for details). IDs for secondary metabolites were omitted, the full metabolic network maps can be found in (Supplementary Figures S5 and S6).

### DNNGIOR-generated models have conservative but precise carbon usage profiles

Finally, we provide an experimental benchmark by comparing the ability of models constructed with DNNGIOR and CarveMe (10) to predict experimentally measured carbon usage profiles of 224 different bacteria (31). We found that although the balanced accuracy scores of DNNGIOR models were similar in range to the CarveMe models, some models performed worse and other models better with no significant trend either way. DNNGIOR’s half-interval algorithm tends to be more conservative in adding reactions than CarveMe’s algorithm, which resulted in more true negatives but also fewer true positives (Figure 8b). Interesting to note are the balanced accuracy scores of W4 DNNGIOR models of Leaf412 and Leaf456, two *Methylophilus* species that showed high scores compared to both the W1 (+0,24) and CarveMe models (+0.34). These high scores are likely due to the low strain versatility, i.e. only glucose and methanol led to in vitro growth, which made an NN-guided accurate inclusion particularly effective.

**Figure 8:**
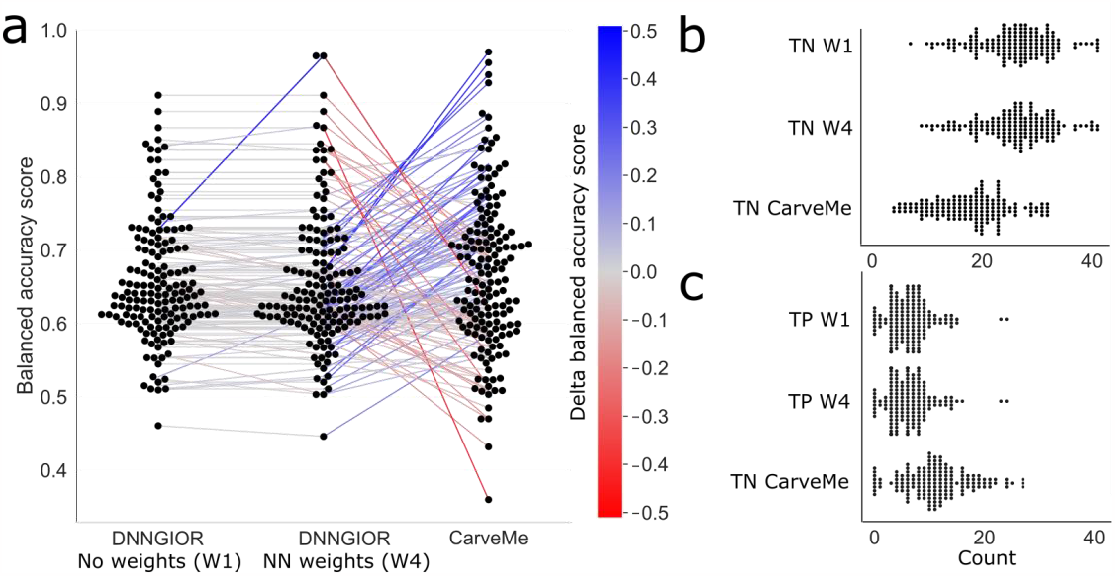
Accuracy of simulated carbon usage profiles. Swarm plots of the a) Balanced accuracy score, b) TNs, and c) TPs of the carbon usage profiles of automatically constructed models compared to experimentally measured profiles. Lines connect scores belonging to the same models showing possible trends. Balanced Accuracy = TPR+TNR/2, TPR = True Positive Rate. TNR = True Negative Rate, TP = True positives, TN = True negatives. For W1 (“No weights”) all reactions in the database are weighted equally. For W4 (“NN weights”) the prediction scores generated by the DNNGIOR neural network were used. Models that showed growth without any carbon source provided or did not grow on any of the carbon sources were omitted. The full carbon utilisation profiles can be found in Supplementary Table 10.

In contrast to the internal validation, using NN weights (W4) did not greatly improve accuracy compared to the gap-filling with no weights (W1). The similarity in scores illustrates that matching phenotypes remains challenging, and manual curation will remain valuable to resolve ambiguities (31). However, DNNGIOR weights provide additional information useful for both manual and automated model reconstruction efforts that represent the evolutionary associations between metabolic reactions throughout the bacterial domain.

### Future potential of the neural networks

The DNNGIOR neural network-based reaction weights significantly improved the gap-filling of GSMMs for a wide variety of bacteria. This success derives from the network learning aspects including the frequency and co-occurrence relationships between reactions. Additional features outside the scope of the current study, such as reaction fluxes, pathway annotation or environmental factors that were previously shown to be useful for gap-filling (23–25,45–47) may also be included, which could improve the prediction accuracy of the weights and subsequent gap-filling even further. A further promising avenue is using the network structure of the input data, including phylogenetic and metabolic networks. Graph convolution takes networks as input and may be a powerful tool to address such complex problems (48). A promising approach using hypergraph link prediction based on the stoichiometric matrix has recently been suggested as an alternative method for gap-filling metabolic networks, showing the potential of including the graph topology into the neural network (24). Combining this with the broader taxonomic signal found in our more diverse dataset could, in the future, lead to improved performance over a broad range of organisms.

### Extensions to increase applicability

Currently, efforts are being made to reconcile the different reaction databases into MetaNetX (49). Once finished, a new neural network could be created that would use the reconciled database and that would be universally applicable. In this paper we focused on ModelSEED models, but we have also created a version trained on CarveMe models showing similar results (Supplementary Figure S8). This shows that it is possible to create neural networks for weighing the reactions in models from different sources. The neural network gives a prediction score for all reactions, not just for the missing ones. Currently, most of these predictions are ignored as we only aimed to fill in the missing reactions and retain the ones that were based on genome annotation, i.e. that have genetic evidence. However, mistakes during metagenome binning (13,14) and annotation do not only result in missing reactions (incomplete MAGs) but also reactions that are falsely attributed to a genome (contaminated MAGs) and thus spurious reactions in the corresponding metabolic models (50). Combining evidence about the predicted taxonomic affiliation of metagenomic contigs (51) with the weights predicted by the DNNGIOR neural network could potentially help identify such erroneous reactions which could then be removed during model curation.

### Final remarks

We developed DNNGIOR, a neural network that predicts which metabolic reactions are present in a given bacterial strain, based on incomplete information. As the neural network learns about these reactions from known bacteria, it is particularly effective for scoring reactions that are relatively common, and for organisms that are relatively closely related to those present in the training dataset. Advanced users can tailor the training dataset for their specific needs, e.g. training models for certain biomes or taxa. The predicted weights can be utilised during gap-filling to improve the accuracy and overall quality of the reconstructed metabolic models. Increasingly, genome-scale metabolic models are being used to interpret microbial metabolic traits, growth, or environmental associations. DNNGIOR should be a valuable tool to enhance the potential of these models.

## Acknowledgements & funding

The authors would like to extend their gratitude to the TBB group for moral support and to J.K. van Amerongen for technical support. Funding support for this work came from the UU-NIOZ project “Turning the tide” (NZ4543.26) awarded to B.E. Dutilh and A.F. Haas, the European Research Council (ERC) Consolidator grant 865694: DiversiPHI, the Deutsche Forschungsgemeinschaft (DFG, German Research Foundation) under Germany’s Excellence Strategy – EXC 2051 – Project-ID 390713860, and the Alexander von Humboldt Foundation in the context of an Alexander von Humboldt-Professorship founded by German Federal Ministry of Education and Research.

## Author Contributions

Writing – Original Draft, M.D.B.; Writing Review & Editing, C.M., A.F.H., D.G., B.E.D.; Visualisation, M.D.B; Software, M.D.B., H.Z., D.G.; Funding Acquisition, A.F.H. and B.E.D.; Supervision, C.M., A.F.H., D.G., B.E.D.

