## Supplementary Tables 6,7,9,10 for "Improving genome-scale metabolic models of incomplete genomes with deep learning"

**Supplementary Table 6:** Mean F1-score for gap-filling draft-models with 30% of reactions removed using different weighting schemes:

| Weights | Mean F1 |
| --- | --- |
| W1: None | 0.015849 |
| W2: Naive binary | 0.115597 |
| W3: Frequency | 0.203976 |
| W4: NN | 0.221615 |

**Supplementary Table 7:** fold change of F1-score and Wilcoxon ranked test for gap-filling draft models with 30% of reactions removed using different weighting schemes.

|  | W1 | W2 | W3 | W4 |
| --- | --- | --- | --- | --- |
| W1: None | x | 7.29 (p = 4*10^-18^) | 12.87 (p = 4*10^-18^) | 13.98 (p = 4*10^-18^) |
| W2: Naive binary |  | x | 1.765 (p = 4*10^-18^) | 1.917 (p = 4*10^-18^) |
| W3: Frequency |  |  | x | 1.086 (p = 3*10^-18^) |
| W4: NN |  |  |  | x |

**Supplementary Table 9:** Percentage decrease of the F1-score of predictions when different phyla are excluded from training

| *Firmicutes* | *Actinobacteria* | *Bacteroidetes* | *Proteobacteria* |
| --- | --- | --- | --- |
| - 0.18 (p = 2.52*10^-66^) | -0.24 (p = 1.86*10^-39^) | -0.43 (p = 1.01*10^-24^) | -0.25 (p = 3.25*10^-125^) |

**Supplementary Table 10:** Mean F1-score for gap-filling using different weighting schemes of 100x iML1515 with 30% of reactions removed.

| Weights | Mean F1 |
| --- | --- |
| W1: None | 0.028002 |
| W2: Naive binary | 0.047483 |
| W3: Frequency | 0.156699 |
| W4: NN | 0.244905 |
