## Supplementary Figures for "Improving genome-scale metabolic models of incomplete genomes with deep learning"

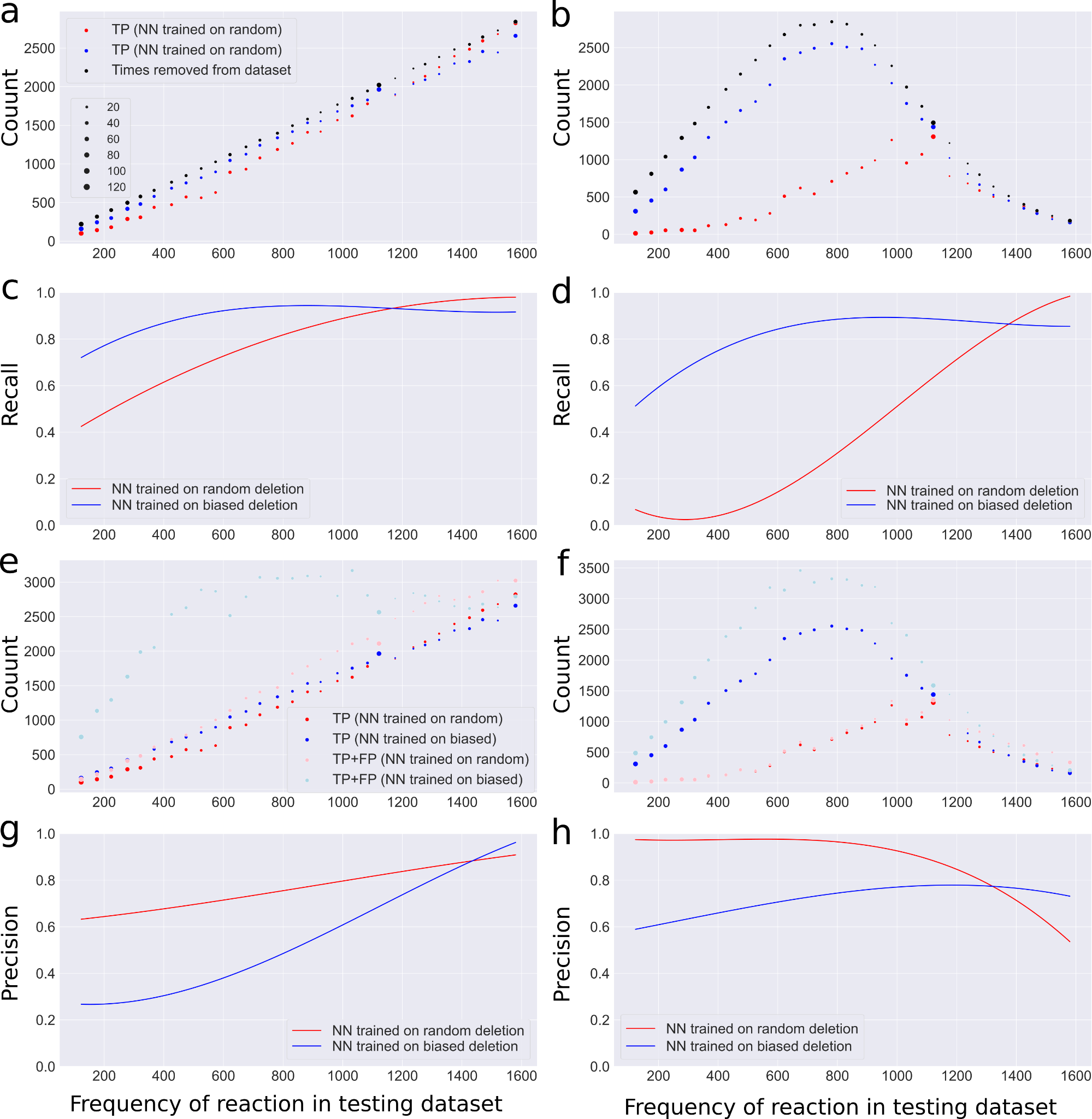

**Supplementary Figure S1 Prediction accuracy on a test dataset with random uniform (a,c,e,g) and biassed deletion (b,d,f,h).** TP = True Positives, FP = False Positives, FN = False Negatives, Recall = TP/(TP+FN), Precision = TP/(TP+FP), NN = Neural Network. a-b) TP counts for both the NN trained on data with random deletion and the NN trained on the training data with biassed deletion and the count for total number of times reactions are removed (TP+FN) for a testing dataset with a) random uniform and b) biassed deletion (see methods). The reactions are averaged over frequency bins of 50 genomes. The dot size corresponds to the number of reactions in the bin. c-d) Polynomial regression curves for the recall of both NNs for a testing dataset with c) random uniform and d) biassed deletion. e-f) Counts of TPs and (TP + FP) for both NNs for a testing dataset with e) random uniform and biassed deletion. Binning of reactions was done as in panel a-b. g-h) Polynomial regression curves of precision of both NNs for a testing dataset with g) random uniform and h) biassed deletion.

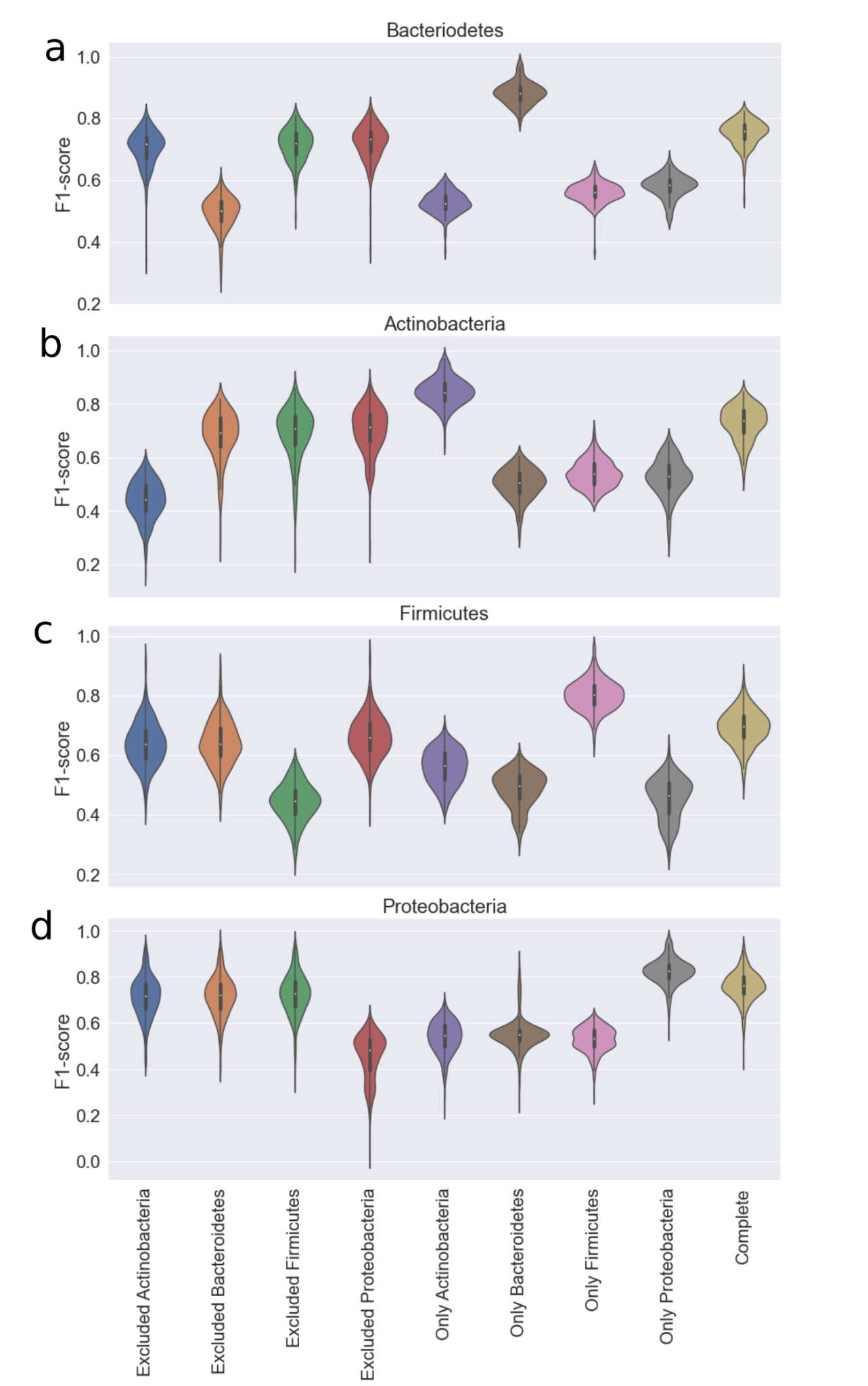

**Supplementary Figure S2:** Violin plots of the prediction accuracy on different when a phylum is excluded from the training data or when the training data contains exclusively one Phylum corrected for training set size for A) Bacteroidetes B) Actinobacteria C) Firmicutes and D) Proteobacteria.

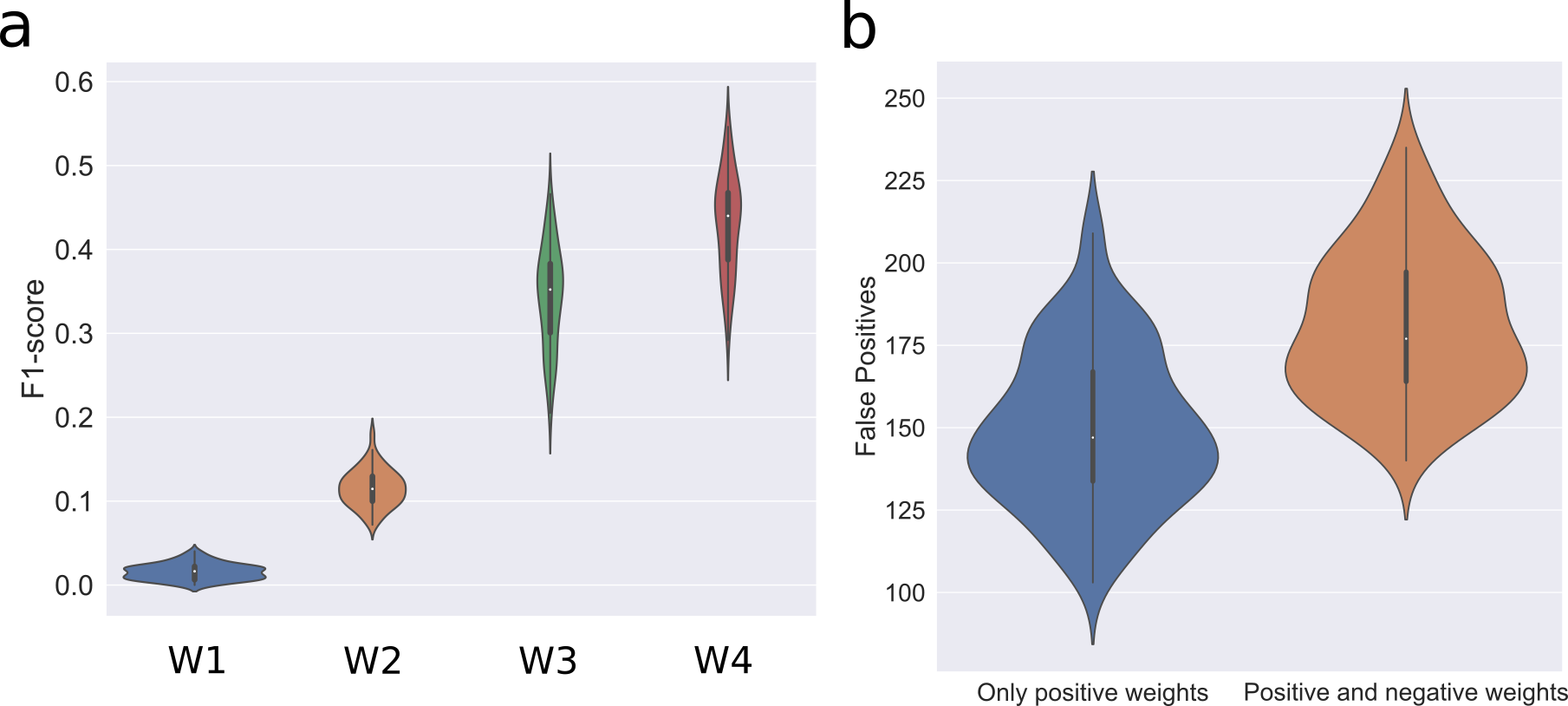

**Supplementary Figure S3: Violin** plots of F1-scores of the gap-filling of 1,659 models in the testing set, from which we randomly deleted 30% of reactions in triplicate. These reduced models were gap-filled with four different weighting schemes where weights ranged from -1 to 1. For W1*: “Naive binary weights” all reactions that are present in the training set were given the same weight of 0. For W3*: “Frequency weights” the frequency of the reaction was used ($W_{freq}= 1- 2*R_{freq}$). For W4* “NN weights” the prediction scores generated by the neural network were used ($W_{NN}= 1- 2*p_{NN}$).

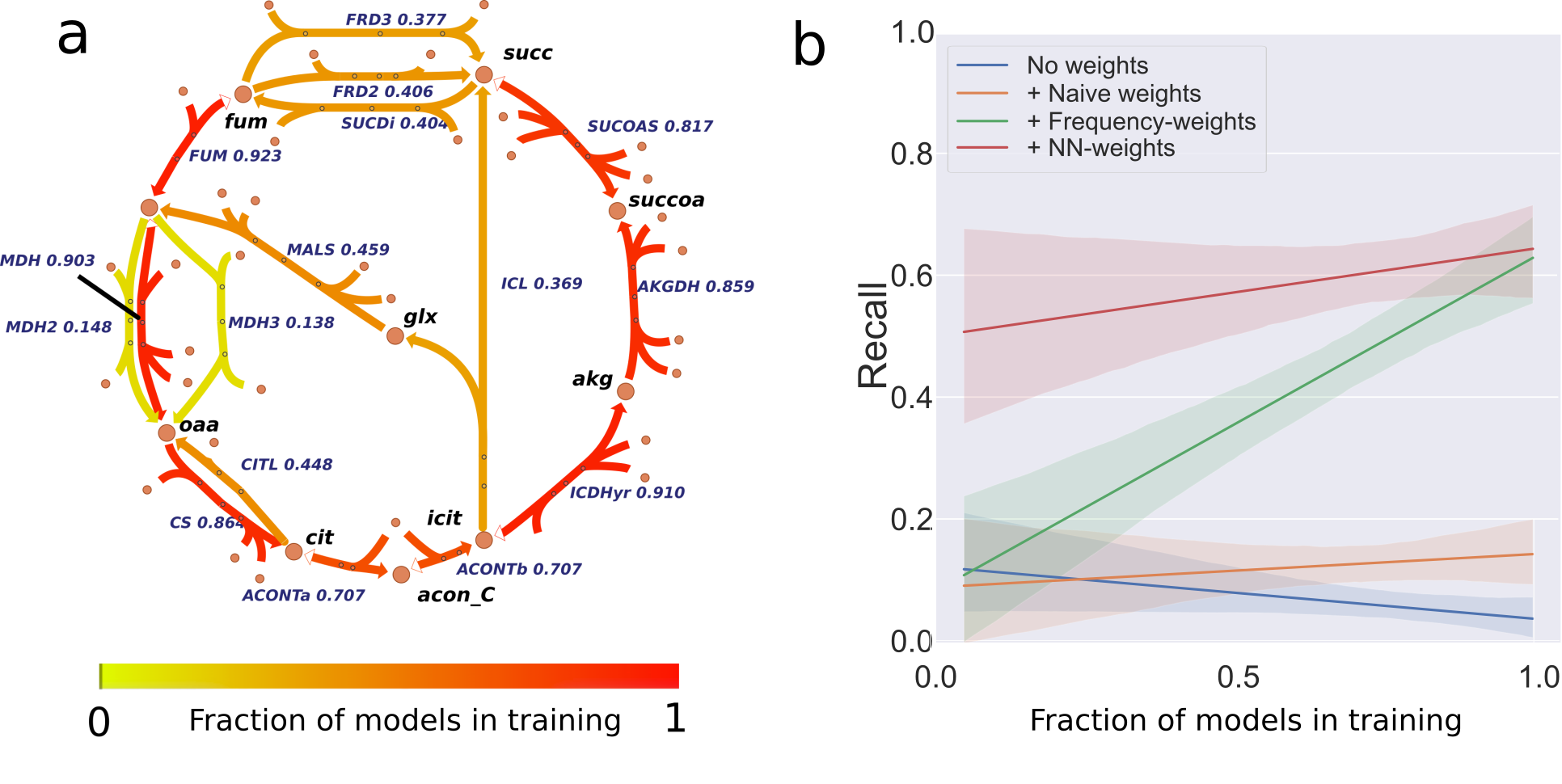

**Supplementary Figure S4: Frequency and recall of reactions in the citric acid cycle.** a) Escher map of the citric acid cycle coloured by frequency. IDs for secondary metabolites are omitted. b) Polynomial regression plots of recall plotted against the fraction of models in training that a reaction is present in for different weighting schemes.

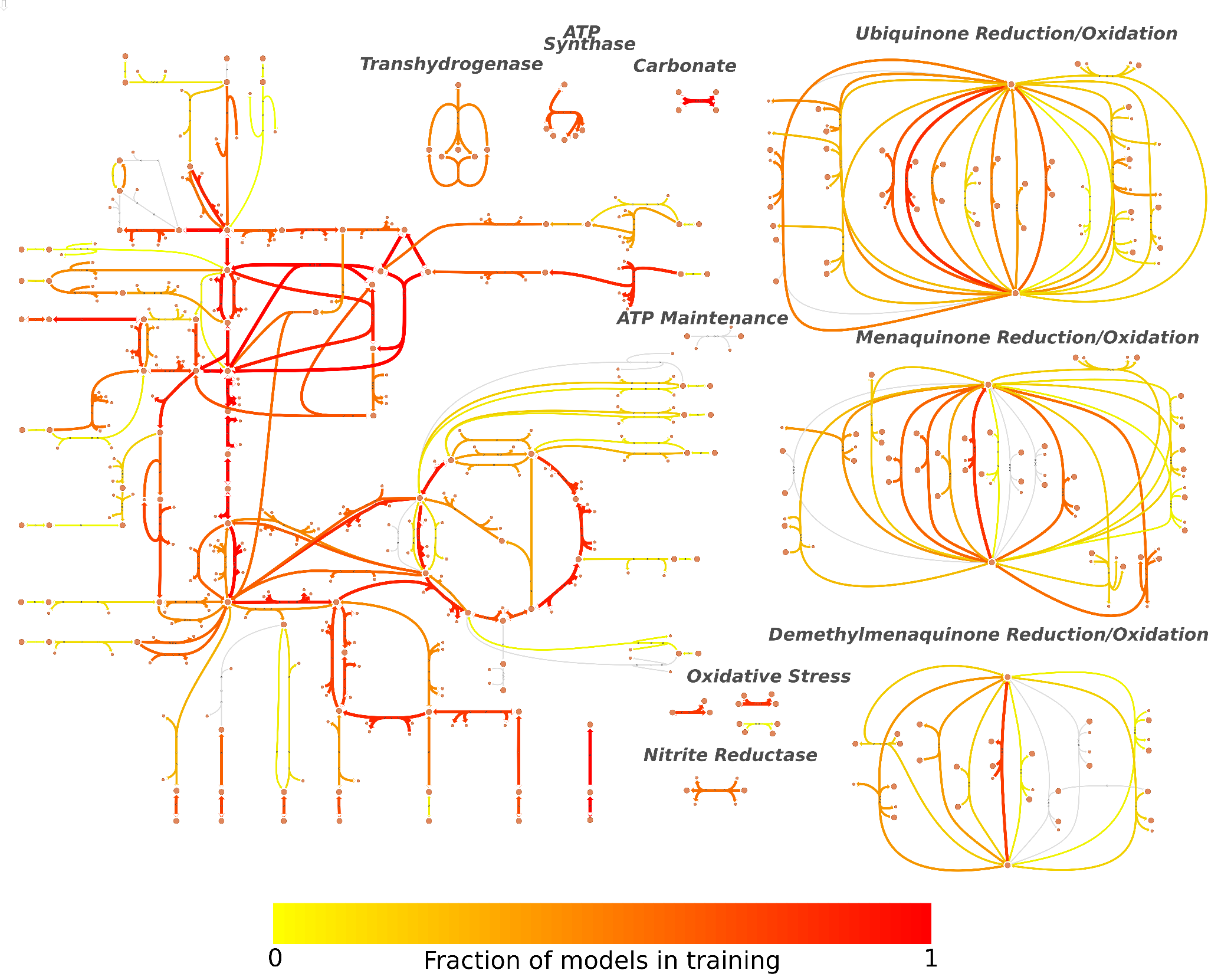

**Supplementary Figure S5: Frequency of the reactions in the central metabolism of *E. coli*.** Escher map of the central metabolism coloured by frequency of the reactions in the models of the training dataset. IDs for secondary metabolites were omitted, reactions in grey were absent from the training dataset.

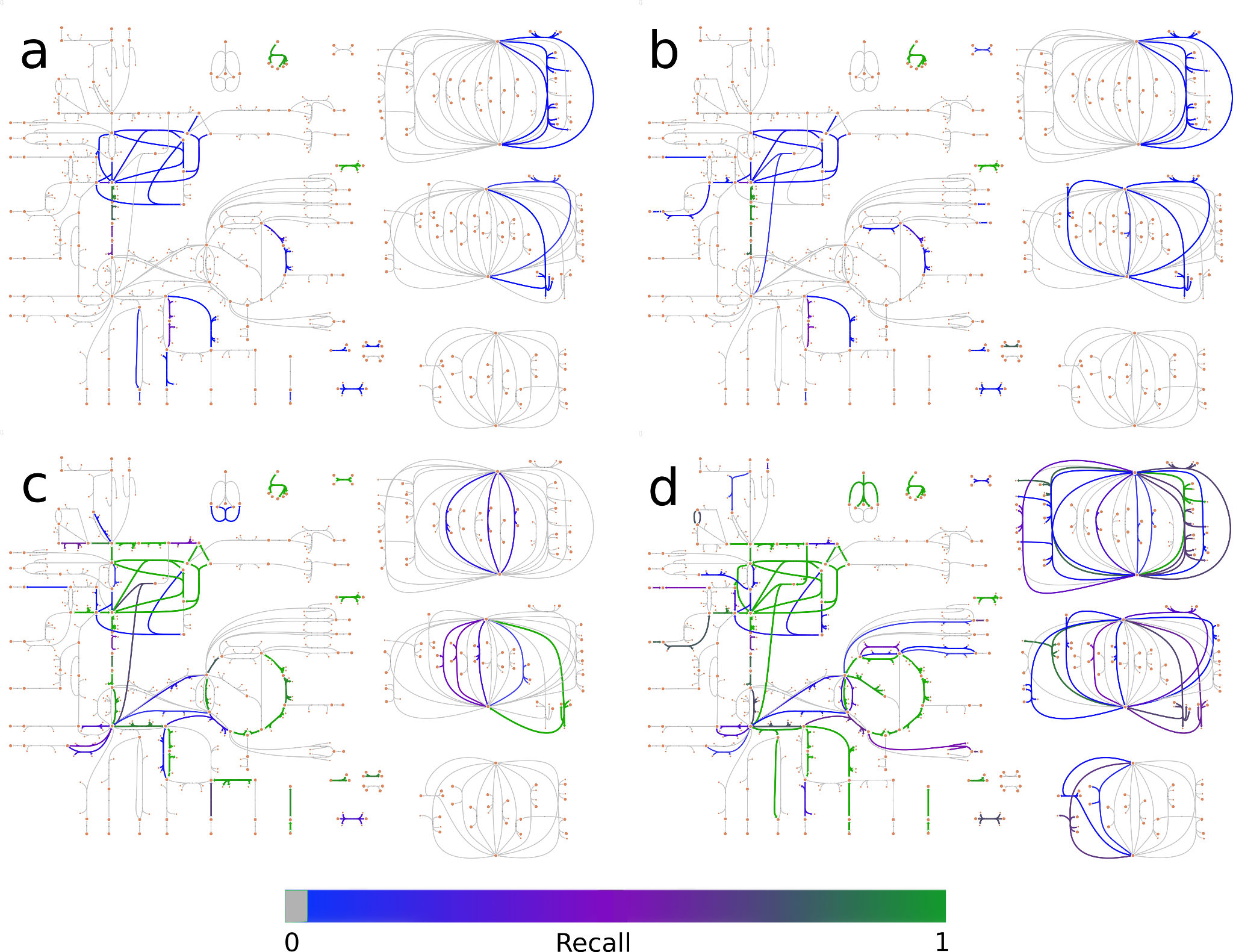

**Supplementary Figure S6: Recall of the reactions in the central metabolism of *E. coli*.** Escher maps of the central metabolism as in Supplementary Figure S6, coloured by the recall after gap-filling models, from which we randomly deleted 30% of the reactions 500 times. Reduced models were gap-filled using four different weighting schemes: a) W1: No weights b) W2: Naive binary weights c) W3: Frequency weights and d) W4: NN weights (see Figure 5 caption and Methods for details). IDs for secondary metabolites were omitted, the full metabolic network maps can be found in (Supplementary Figures S4 and S5).

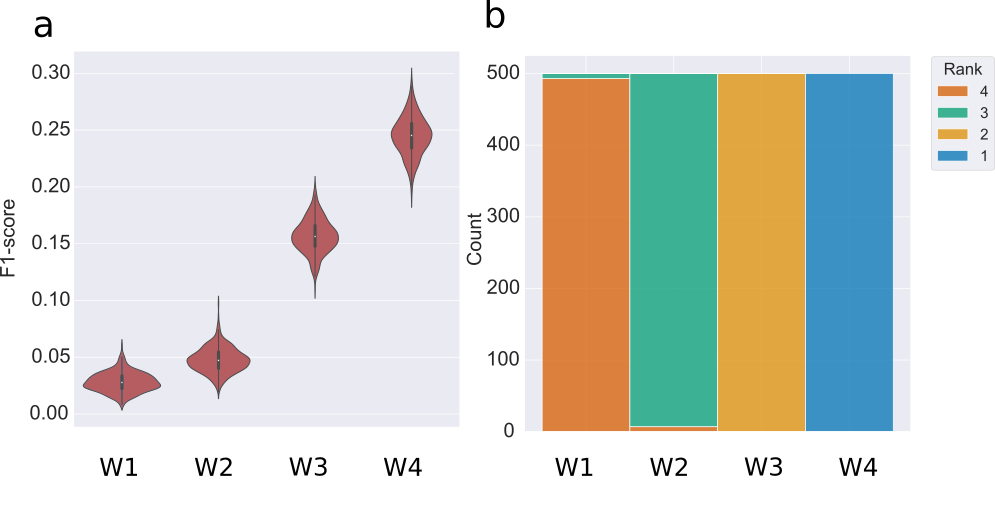

**Supplementary Figure S7:** a) Violin plot of F1-scores of the gap-filling of iML1515 (*E. coli*), from which we randomly deleted 30% of reactions 500 times. These reduced models were gap-filled using four different weighting schemes (Equation 3). For W1 (“No weights”) all reactions in the database are weighted equally. For W2 (“Naive binary weights”) all reactions that are present in the training data were given the same weights. For W3 (“Frequency weights”) the frequency of the reaction was used to weigh reactions. For W4 (“NN weights”) the prediction scores generated by the DNNGIOR neural network were used. b) Distribution of the ranks of the F1-scores for different weights with four being the worst ranking and one being the best.

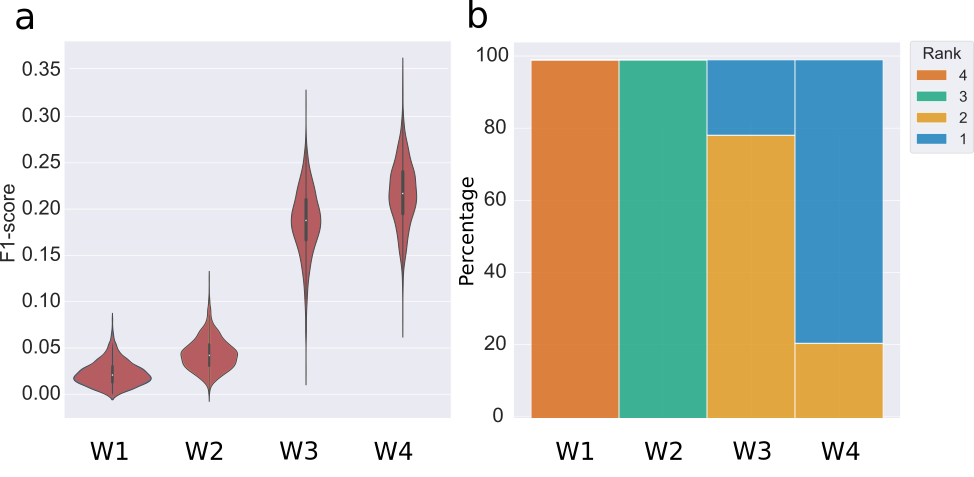

**Supplementary Figure S8**: **Weighted gap-filling of CarveMe models.** a) Violin plot of F1-scores of the gap-filling of 1,659 models from the testing dataset, from which we randomly deleted 30% of reactions in triplicate. These reduced models were gap-filled using four different weighting schemes (Equation 3). For W1 (“No weights”) all reactions in the database are weighted equally. For W2 (“Naive binary weights”) all reactions that are present in the training data were given the same weights. For W3 (“Frequency weights”) the frequency of the reaction was used to weigh reactions. For W4 (“NN weights”) the prediction scores generated by the DNNGIOR neural network were used. b) Distribution of the ranks of the F1-scores for different weights with four being the worst ranking and one being the best.

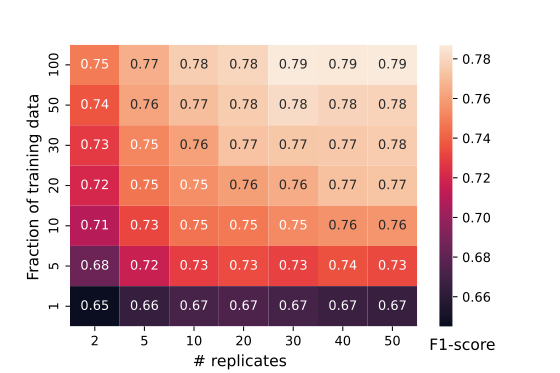

**Supplementary Figure S9: Effect of size training data:** Heatmap of F1-scores of predictions by a neural network with different sizes of training set. Fraction of training data refers to the fraction of the total 13,359 genomes using for training, # replicates is the number of times 30% of reactions were removed to generate features.

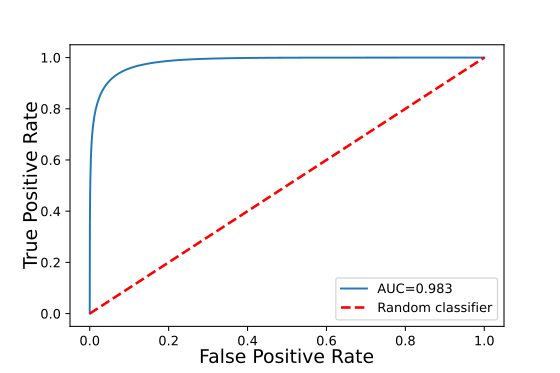

**Supplementary Figure S10 ROC-curve:** ROC plot of predictions made by the neural network on the one-per-genus testing dataset, AUC = area under the curve. The dotted line corresponds to a random classifier.

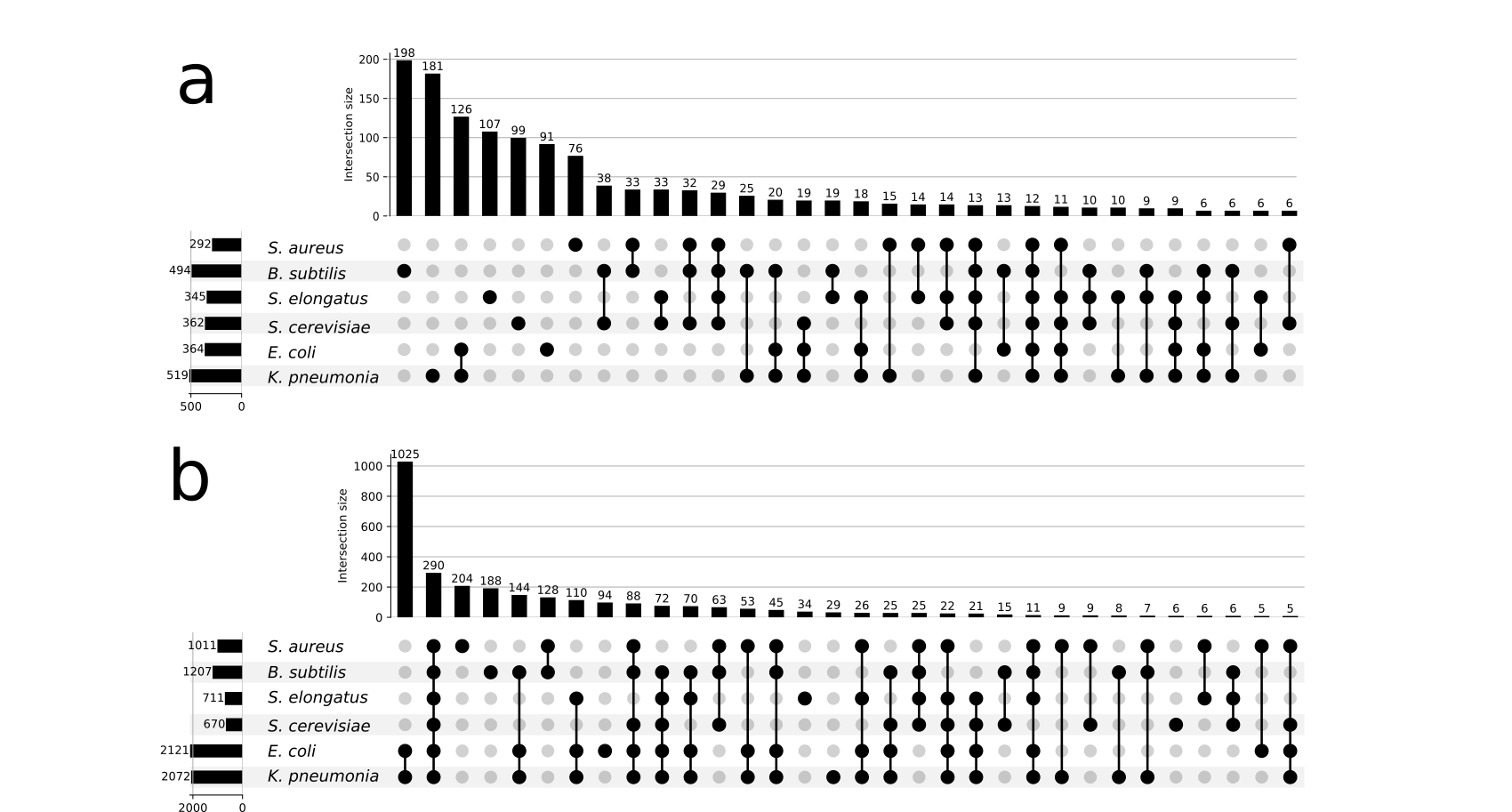

**Supplementary** **Figure S11 DNNGIOR neural network overlap:** upset plots showing the overlap in: a) predicted missing reactions and b) all predictions made for 6 manually curated models, unions with size < 5 are omitted.

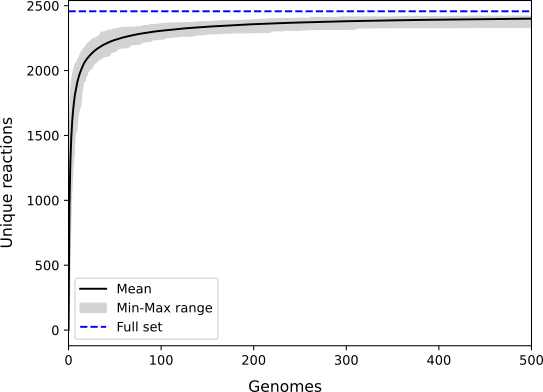

**Supplementary Figure S12 Rarefaction curve of the reactome:** Simulated growth of reactome by sampling of 20 times 500 genomes from the training dataset. The dotted line corresponds to the reactome size of the full set of 13,359 genomes.

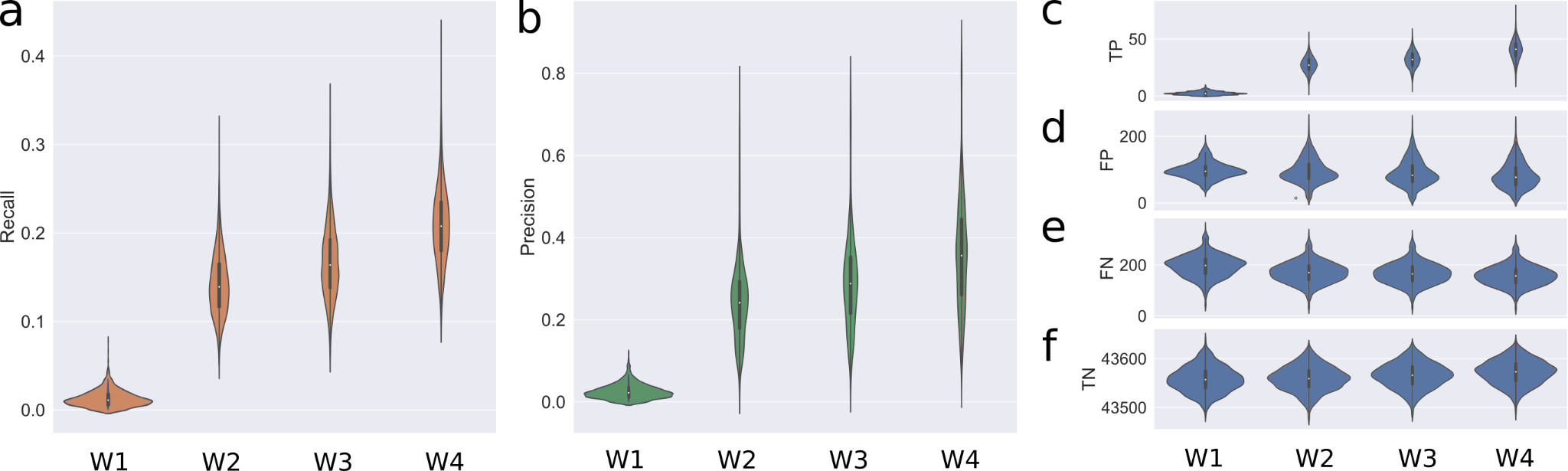

**Supplementary Figure S13 Weighted gap-filling of draft models.** Violin plots of a) Recall-scores, b) Precision scores, c) TPs, d) FPs, e) FNs and f) TNs of the gap-filling of 1,659 models from the testing dataset, from which we randomly deleted 30% of reactions in triplicate. These reduced models were gap-filled using four different weighting schemes (Equation 3). For W1 (“No weights”) all reactions in the database are weighted equally. For W2 (“Naive binary weights”) all reactions that are present in the training data were given the same weights. For W3 (“Frequency weights”) the frequency of the reaction was used to weigh reactions. For W4 (“NN weights”) the prediction scores generated by the DNNGIOR neural network were used.
